# Flexible, Cluster-Based Analysis of the Electronic Medical Record of Sepsis with Composite Mixture Models

**DOI:** 10.1101/160465

**Authors:** Michael B. Mayhew, Brenden K. Petersen, Ana Paula Sales, John D. Greene, Vincent X. Liu, Todd S. Wasson

## Abstract

The widespread adoption of electronic medical records (EMRs) in healthcare has provided vast new amounts of data for statistical machine learning researchers in their efforts to model and predict patient health status, potentially enabling novel advances in treatment. In the case of sepsis, a debilitating, dysregulated host response to infection, extracting subtle, uncataloged clinical phenotypes from the EMR with statistical machine learning methods has the potential to impact patient diagnosis and treatment early in the course of their hospitalization. However, there are significant barriers that must be overcome to extract these insights from EMR data. First, EMR datasets consist of both static and dynamic observations of discrete and continuous-valued variables, many of which may be missing, precluding the application of standard multivariate analysis techniques. Second, clinical populations observed via EMRs and relevant to the study and management of conditions like sepsis are often heterogeneous; properly accounting for this heterogeneity is critical. Here, we describe an unsupervised, probabilistic framework called a composite mixture model that can simultaneously accommodate the wide variety of observations frequently observed in EMR datasets, characterize heterogeneous clinical populations, and handle missing observations. We demonstrate the efficacy of our approach on a large-scale sepsis cohort, developing novel techniques built on our model-based clusters to track patient mortality risk over time and identify physiological trends and distinct subgroups of the dataset associated with elevated risk of mortality during hospitalization.

**Abbreviations:**

EMR: electronic medical record
CMM: composite mixture model
KPNC: Kaiser Permanente Northern California
BIC: Bayesian information criterion
AIC: Akaike information criterion
PAM: partitioning around medoids
MICE: multivariate imputation using chained equations

## 1. Introduction

Electronic medical records (EMRs) have become increasingly ubiquitous in healthcare, and the utility of these complex datasets for clinical decision support is the subject of much current research ([7] and references therein). EMR data comprise multivariate observations of variables with discrete or continuous values. These variables can be static (observed only a single time during a hospitalization; e.g. gender) or dynamic (e.g. vital signs). Deriving actionable insights from EMR data requires appropriate models for these multi-typed observations [29].

Besides complexity in the types of information contained in the EMR, heterogeneity inherent in the clinical population under study adds to the challenge of modeling these data. This physiological heterogeneity is a hallmark of debilitating conditions like cancer [16] and, in particular, sepsis [25], an increasingly prevalent clinical condition characterized by a dysregulated immune response to infection leading to organ dysfunction and death. Accounting for this heterogeneity can have considerable therapeutic importance. In the case of breast cancer, for example, stratification of patient tumors into molecular subtypes significantly increased precision of treatment and improved survival (reviewed in [34]). In the case of sepsis, delays in antibiotic administration lead to considerably elevated mortality risk [23], suggesting that a technique highlighting subtle physiological phenotypes associated with elevated mortality risk early in an inpatient stay might aid triage and treatment of potentially septic patients. Indeed, a central goal of this and other similar efforts is to identify physiologically distinct subgroups of a clinical population more at risk for adverse health outcomes that can be targeted with interventions tailored to the subgroup’s characteristics [22, 39, 30, 15, 19, 26, 40].

In this study, we describe a joint probabilistic framework called a composite mixture model (CMM; [32, 37]), a technique heretofore never applied to EMR data. The CMM accommodates the wide variety of data types common in EMR datasets while accounting for heterogeneity in the clinical population. We adapt our model to analyze a large EMR dataset composed of more than 53,000 emergency department (ED) hospitalization episodes from Kaiser Permanente Northern California (KPNC) in which patients were suspected to have infection and, in a subset of cases, met the criteria for sepsis [36, 8]. We demonstrate the efficacy of our data-driven approach by identifying and annotating clusters of patient episodes with significantly higher risk for mortality during hospitalization, by visualizing physiological trends associated with these elevated rates of mortality, and by benchmarking the performance of our framework on common EMR analysis tasks such as missing data imputation.

## 2. Related Work

One approach to modeling EMR data focuses on the important task of prediction in order to prevent or mitigate an impending adverse outcome. Traditional predictive modeling approaches, such as logistic regression and random forests [5], can provide useful insights for clinical decision support. For example, multi-task predictive models have proven successful in stratifying patients according to risk of developing hospital-acquired infections [39, 40]. Other approaches have been able to recover clinically relevant phenotypes with striking prediction capabilities for a wide range of medical conditions [15, 19]. Particularly in sepsis, supervised approaches have demonstrated remarkable performance in early identification of patients at risk of entering septic shock [18, 17]. In essence, these models characterize the effects of a fixed set of predictor variables or features on an outcome of interest (*supervised* learning; a *conditional model*), without directly modeling the features themselves. These approaches also require complete case data (i.e. observations without missing entries). Without first performing imputation, certain samples or even entire features could be removed from analysis. More importantly, many of these approaches assume that the population under study can be modeled as a single, homogeneous group, ignoring the possibility that the population represents a mixture of distinct subgroups potentially amenable to different treatment regimes. In contrast, we develop a framework able to account for both population heterogeneity and missing data and, in an unbiased manner, aim to identify latent phenotypes in the population enriched for mortality events during hospitalization rather than predict mortality events or cataloged conditions (e.g. ICD9 codes).

Unsupervised approaches, on the other hand, learn structure or dependencies by modeling the whole data observation (outcome and features) together, capturing dependencies among the different dimensions of the full observation vector. Such approaches have proven successful recently in directly modeling complex EMR observations or embedding them in lower-dimensional feature spaces that can then be used for prediction or risk stratification [22, 30, 2, 26]. Multivariate statistical mixture models are another class of unsupervised learning approach that can formally represent heterogeneity in the population, treating both outcomes and features as probabilistic quantities to be modeled jointly (*joint* model; [33]). However, such mixture models are difficult to apply in their conventional form when elements of a multivariate observation are of different types (e.g. a categorical variable like gender and a continuous variable like median diastolic blood pressure) as is common with EMR data. Similar unsupervised work has recently been presented to model these data (e.g. [22] and [30]). However, these studies either focused on a single clinical variable (uric acid trajectories [22]) or transformed multi-typed EMR observations into a single representation for modeling purposes (e.g. bag-of-word representations [30]). In contrast, we directly model 32 different physiological and demographic features of both discrete and continuous types with our simple and extensible joint probabilistic framework.

## 3. Methods

### 3.1. KPNC Sepsis Cohort Description

Kaiser Permanente Northern California is a highly integrated healthcare delivery system with 21 medical centers caring for an overall population of 4 million members. The full KPNC dataset consists of 244,248 in-patient hospitalization visits (mortality rate: 5.2%) with a suspected or confirmed infection and sepsis diagnosis, drawn from KPNC medical centers between 2009 and 2013 [36]. We refer to each in-patient visit as an *episode* for the remainder of this work. For this analysis, we used a subset of the full dataset consisting of 53,659 in-patient episodes, with an overall mortality rate of ∼6%. Our criteria for creating this analysis cohort were the following: 1) hospitalization admission occurred via the emergency department (to identify high-risk patients earlier in their care delivery); 2) the length of hospitalization was at least twelve hours; and 3) all vital signs were taken three or more times during the first three hours of hospitalization. Figure 1 illustrates the process by which we prepared our different analysis cohorts.

**Figure 1:**
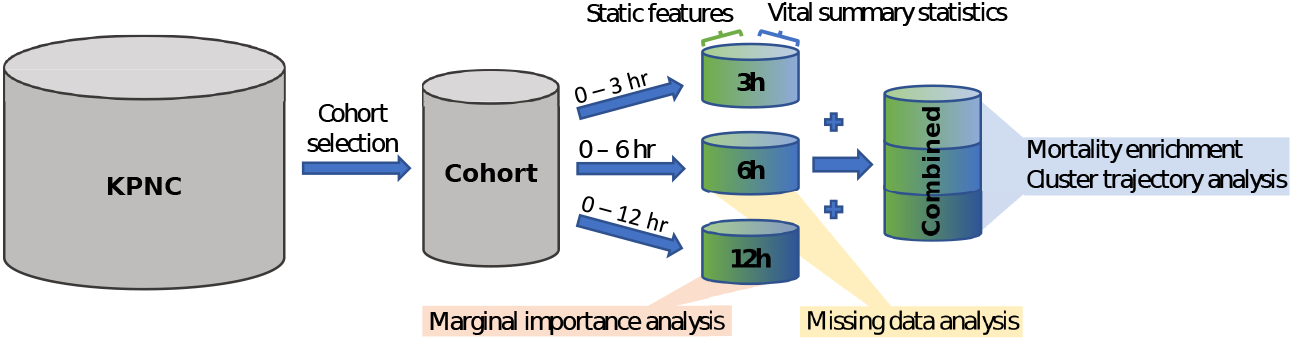
Flowchart of cohort selection, data pre-processing, and analysis dataset construction.

Our cohort consisted of both static and dynamic features. The static features that made up our dataset include: 1) age; 2) sex; 3) treatment facility code; 4) KPNC membership status; 5) indicator of whether patient was transported in from another site; 6) LAPS2, a KPNC single measure of *acute* disease burden at the time of hospital admission; 7) COPS2, a monthly KPNC aggregate measure of *chronic* disease burden; and 8) an indicator of patient mortality status at the end of the episode. The vital signs included in this analysis were 1) diastolic blood pressure, 2) systolic blood pressure, 3) heart rate, 4) respiratory rate, 5) temperature, and 6) pulse pressure. Outlying vital sign observations were removed according to the following filters: heart rate > 300; systolic blood pressure > 300; respiratory rate > 80; body temperature > 108; and body temperature < 85. We then computed the maximum, minimum, median, and standard deviation of patient vital signs over different post-admission periods (3, 6, or 12 hours). Pairing the summary statistics with their corresponding static admission features resulted in three analysis cohorts, one for each postadmission period (Figure 1).

### 3.2. Composite Mixture Model Definition

Here, we describe the composite mixture model, a flexible joint probability model for multi-typed, multivariate data. The two central ideas behind the CMM are that: 1) the population is heterogeneous (composed of subgroups or clusters) and 2) we can specify the full joint distribution of a multi-typed observation vector, x, by specifying appropriate univariate, exponential family distributions for each dimension of x. The CMM takes the following form:

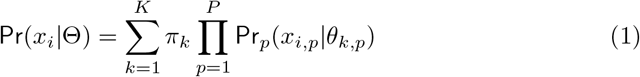

where *x_i_* is an observation vector of dimensionality *P, K* is the number of mixture components or clusters in the model, Pr*_p_* is the distribution of the *p*^th^ dimension (e.g. univariate Gaussian), and *θ_k,p_* are the model parameters for the *p*^th^ dimension distribution in the *k*^th^ mixture component. This structure is reflected in the plate notation diagram in Figure 2. For completeness, we show an equivalent formulation of the CMM model in Figure 2 that includes the indicator variables, *Z_i_*, categorical variables that take the value *k* (from 1 to *K*) if episode *i* is assigned to cluster*k*. For example, suppose we had a three-dimensional observation vector wherein the first dimension contained integer count data, the second dimension contained positive and negative real values, and the third dimension contained discrete categorical data. A potential CMM for these data might consist of the following univariate distributions with different parameter values for each distribution depending on the mixture component *k* to which *x_i_* was assigned: 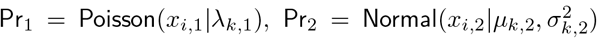, and Pr_3_ = Categorical(*x_i_,3 |γ_k,3_*) (*γ_k,3_* is the vector of probabilities for each possible discrete value of *x_i,3_*). The model structure implies that, given a mixture component, the dimensions of the observation vector are independent of one another. It is important to note that, despite the features being treated as independent, complex correlations among features can be recovered by mixing (or averaging) over more components.

**Figure 2:**
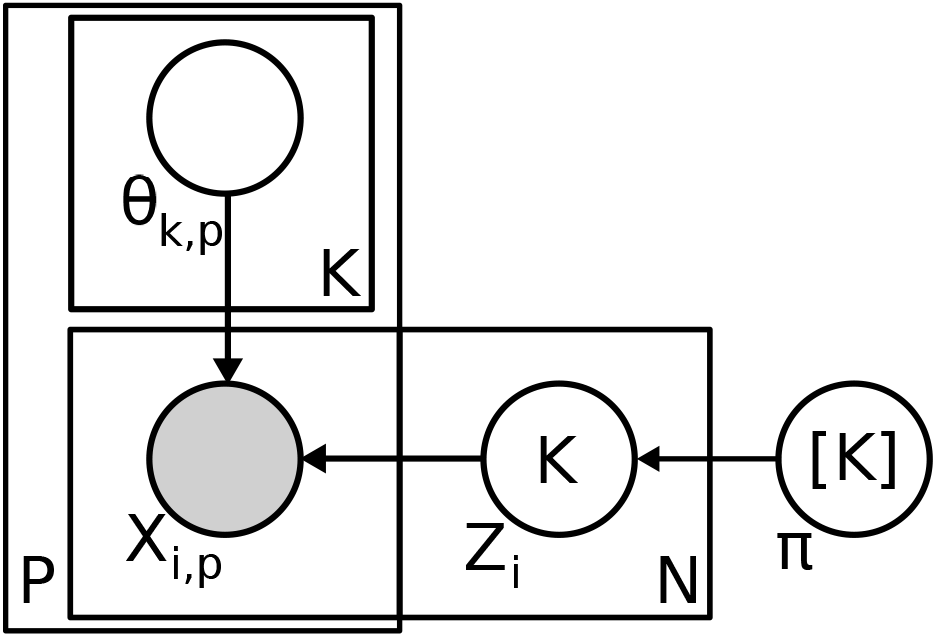
Plate notation diagram of composite mixture model. To indicate the relationship between episodes and clusters, the formulation shown here includes the latent indicator variables, *Z_i_*, that specify the cluster assignment for episode *i*. This formulation is equivalent to the one shown in equation 1.

### 3.3. Composite Mixture Model Fitting to KPNC Dataset

As shown in Section 3.2, fitting a composite mixture model first requires specification of the univariate distributions for each dimension of an observation vector. We support exponential family distributions for each observation dimension that can be efficiently estimated in parallel by computation of sufficient statistics. For our analysis, categorical variables (including mortality status) were modeled with categorical distributions while age, LAPS2, and COPS2, and the summary statistics of patient vital signs were modeled with either normal, gamma, or exponential distributions where appropriate. To facilitate model estimation, we added a small amount of jitter to non-categorical features. Specifically, we added a uniform random sample in the range −*d*/10 to *d*/10 to each value, where d is the smallest distance between unique values of that feature.

To fit a model for a given number of clusters, we performed expectation-maximization [9]. As expectation-maximization is a local optimization approach sensitive to initialization, we performed 3 random restarts of each fitting procedure with a maximum of 100 iterations (we observed that this number of iterations was generally sufficient for adequate model fitting). We determined the optimal number of clusters by evaluating the Bayesian information criterion (BIC) of all fitting restarts for each number of clusters tested (from 10 to 1000 clusters in increments of 10 for the individual post-admission datasets) and selecting the model with the lowest BIC. We also considered alternative model selection criteria such as Akaike information criterion (AIC), though BIC generated more reasonable results considering its harsher penalty on model complexity. All composite mixture model data structures and fitting routines were implemented in a custom R software package (available upon request).

### 3.4. Mortality Enrichment and Cluster Trajectory Analysis

To identify changes in the physiological status of a potentially septic patient during hospitalization, we combined the 3h, 6h, and 12h datasets into a single dataset (hereafter referred to as the *combined* dataset) and fit a CMM to all observations (Figure 1). In this way, we treat the three observations for each episode as independent, and while the static features were shared (driving all three observations of a single episode to co-cluster with one another), changes in the vitals over the course of hospitalization drive association of a given episode with different clusters over time. The first motivation for this choice was to characterize latent phenotypes across the whole clinical population and every post-admission period. Since patients would likely enter the emergency department at different stages of their condition, this time-agnostic clustering would allow a late-stage, debilitated patient within the first 3 hours after admission to be co-clustered with a patient who reached a similar physiological state at, say, 12 hours post-admission. The second motivation for this approach was to address the label-switching problem [35], a common challenge in mixure modeling. Fitting each dataset separately would have resulted in different sets of labels for each post-admission period that could not have been directly linked over time (i.e. cluster 1 at 3 hours post-admission would likely not have corresponded to cluster 1 at 6 hours post-admission).

Consequently, to generate cluster assignments for each episode and postadmission period, we fit a CMM to the combined dataset, resulting in an optimal number of 650 clusters (tested 50 to 1000 in increments of 50), with cluster sizes ranging from 88 to 2,883 observations. As this number of clusters was prohibitively large for visualization purposes, we then clustered the estimated parameters of the fitted model using the partitioning around medoids (PAM) algorithm [20] with the final number of clusters set to 20. These cluster labels were used in analyses for Figures 4, 5, and 6. We then mapped the newly assigned cluster labels to each observation in the dataset.

Once we assigned cluster labels to each combination of episode and postadmission period, we combined the cluster labels for each of an episode’s three post-admission periods into a single vector (*cluster trajectory*). As with our annotation of mortality enrichment for each post-admission period, we computed the significance of enrichment of each cluster trajectory for mortality events using a one-tailed Fisher’s exact test. We adjusted for multiple hypothesis testing with a Bonferroni-corrected significance level of ∼ 2.51e-05. (Figure 5).

To evaluate the physiological signatures of the three clusters at 12h postadmission in the top mortality-enriched cluster trajectories, we first normalized the LAPS2, COPS2, and vital sign features to values between 0 and 1 by performing feature scaling (i.e. subtracting each feature by its minimum and dividing by the difference between the feature’s maximum and minimum). We then conducted Wilcoxon rank-sum tests, comparing each cluster-specific distribution of LAPS2, COPS2, and vital sign statistics with their respective distributions in the overall population and computing 95% confidence intervals on the estimated difference between the cluster and overall population (Figure 6).

### 3.5. CMM-Derived Marginal Importance Analysis

Marginal importance plots are often used in conjunction with random forest models to visually evaluate the marginal effect of different features on the rate of a particular outcome [5, 21]. We adapted this concept to develop novel visualizations based on CMM clusters. We first determined the optimal number of clusters (200) for the 12h post-admission dataset (chosen arbitrarily) using the BIC criterion as described in Section 3.3. Unlike in the previous analyses with the combined dataset, we retained the estimated parameters for this model 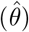 instead of clustering them further using PAM. The procedure we followed to compute the two essential components of our CMM-based marginal importance plot (estimated mortality rates 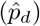 and estimated sample sizes 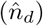 at different values of a vital sign feature of interest) is shown in Figure 3. Briefly, from a dataset *X* (*N* episodes, *P* features), we compute the matrix, *W*, containing the posterior probabilities of assigment of each episode to one of the K clusters (each row of *W* sums to 1). If we consider the sample size of a single episode to be 1, these probabilities represent the distribution of that sample size over the *K* clusters. Simultaneously, we generate a dummy feature matrix for a feature of interest *f* (lower left of Figure 3). In this dummy feature matrix, we enumerate a column containing a range (from their lowest to highest observed values) for a feature of interest. All other feature columns are set to 0. We then use the estimated CMM model parameters, 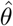, to compute the matrix, *M_f_*, containing the posterior probabilities of assignment of each dummy feature row to one of *K* clusters. We also compute the expected rate of mortality 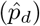 given each value in the range of *f*, by averaging the estimated conditional frequency of mortality in each cluster over all *K* clusters. Finally, we compute the estimated sample sizes 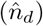 at each value of *f* by multiplying the *W* by the transpose of *M* and taking the sum of the rows of the resulting *N X F* matrix. In this way, we computed the cumulative density (the estimated number of episodes) across all episodes represented at each value in the range of a feature of interest. The range of feature values becomes the domain of our marginal importance plot and mortality rate is plotted on the y-axis, with the estimated mortality rate 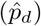 and sample sizes 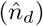 used to construct the curve and 95% Wilson score confidence intervals ([41]; shown in Figures 7a, 7b, 7c, 7d). We note that, even after filtering patient vital signs for outliers, certain extreme values can still be observed during a typical hospitalization ([10]; e.g. blood pressure values of 0 for cardiac arrests; Figure 7a).

**Figure 3:**
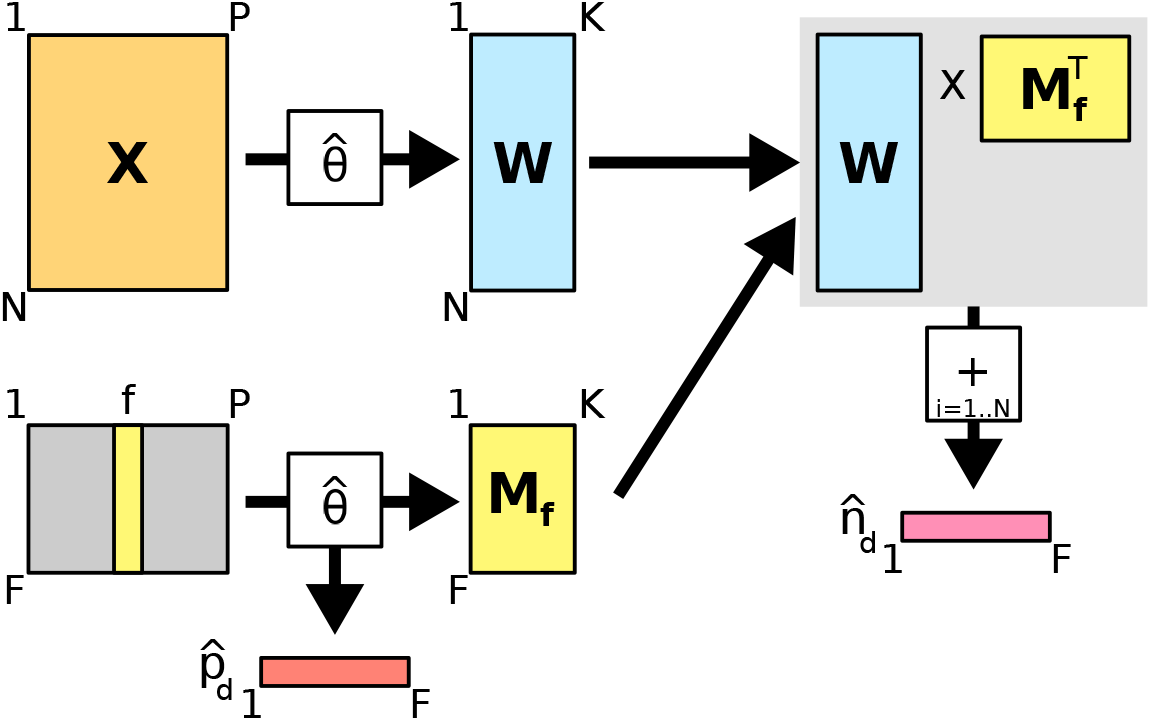
Diagram of computational procedure for constructing CMM-based marginal importance plots.

### 3.6. CMM-Based Missing Data Imputation

For our missing data analysis, we introduced missing entries (completely at random; MCAR) at a rate of 5% in the 6 hr dataset (arbitrarily chosen) for all features except age, gender, and mortality, which were assumed non-missing for all records. The resulting dataset contained 12,344 complete records (i.e. records with no missing values).

Imputation using the CMM involves three steps: 1) performing a CMM fit on either A) the remaining complete records of the MCAR dataset or B) the MCAR dataset after performing population mean imputation, 2) re-computing the cluster membership probabilities for each episode conditioned on the non-missing features, and 3) imputing each missing value with either A) the cluster-averaged expected value (for non-categorical variables) or B) the label associated with the mode of the cluster-averaged expected category probabilities (for categorical features).

We compared imputation using the CMM to population mean imputation (as a baseline), multivariate imputation using chained equations (MICE) [6, 38], and *k*-nearest neighbors imputation. For our purposes, we set the prediction method of MICE to predictive mean matching [6] for non-categorical variables and logistic/polytomous regression for categorical variables. For *k*-nearest neighbors, we tested neighborhoods of size 5 and 20, assigning the value with the largest frequency (’majority vote’) among neighbors for discrete, categorical variables or the mean of neighbor values for continuous variables.

For each feature column with missing values, imputation performance was characterized by a distance, *D*. For non-categorical features, we used the sum of squared deviations from the observed values: 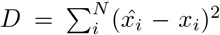, where *x_i_* and 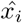 are the observed and imputed values for the *i*^th^ imputed record, respectively, and *N* is the total number of imputed records. For categorical features, we used the counts of incorrectly imputed values: 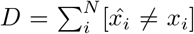, where brackets indicate 1 when the condition is true and 0 otherwise. Distances for each feature were then normalized by the corresponding distance obtained using population mean imputation. Rankings of distance across imputation methods were robust to the choice of distance metric (squared deviations or absolute deviations for non-categorical features; overlap, occurrence frequency, inverse occurrence frequency, Eskin [11], Goodall [13] for categorical features).

## 4. Results

### 4.1. CMMs Reveal Temporal & Physiological Signatures of Mortality Risk in Sepsis

An important challenge in treatment of sepsis is the detection of (sometimes subtle) physiological changes in a patient while accounting for the heterogeneous presentation of the condition. We characterize these latent changes by modeling the full demographic and physiological space of all episodes, regardless of time, subsequently identifying the regions or clusters of this space with which a patient episode associated during an ED hospitalization. To this end, we concatenated the 3h, 6h, and 12h datasets into a *combined* dataset, fit a CMM to all observations, and generated assignments for each episode at each post-admission time to one of twenty clusters (see Figure 1). This approach allowed us to characterize demographic and physiological heterogeneity across all episodes regardless of time and severity of condition (more discussion of rationale in Methods section 3.4). Despite our use of vital sign summary statistics as features, we observed considerable feature variability within each of the 20 clusters, indicating that clustered episodes were not assuming single, repeated values for any feature. We ascribe clinically relevant annotations to our clusters by assessing each of them for statistical significance of enrichment for mortality events during hospitalization. We note that this data-driven cluster annotation method could just as easily been applied to other binary-valued adverse health events (e.g. ICU transfer, need for mechanical ventilation, etc.), indicating the portability of our approach to other settings or healthcare centers.

From this analysis, we determined the proportion of episodes in each cluster at each post-admission time and identified those clusters more or less associated with mortality events. We found that while the majority of the episodes were assigned to one of the top seven clusters over all post-admission periods, some clusters were associated with progressively fewer episodes from 3h to 12h post-admission (e.g. cluster 8) while other clusters (e.g. cluster 10) became associated with more episodes over time (Figure 4a). These patterns highlight the diverse demographic/physiological “landscape” through which patients progressed during their hospitalization and indicate how some clusters represent transient “stops” in a patient’s progression. When we considered each cluster separately, we found that mortality events were significantly underrepresented in episodes assigned to clusters 2, 4, and 5. Conversely, clusters 1, 3, 10, 13, 14, 15 and 17 showed high degrees of enrichment for mortality events (Figure 4b), indicating population heterogeneity even among those patient episodes considered at high risk for mortality.

**Figure 4:**
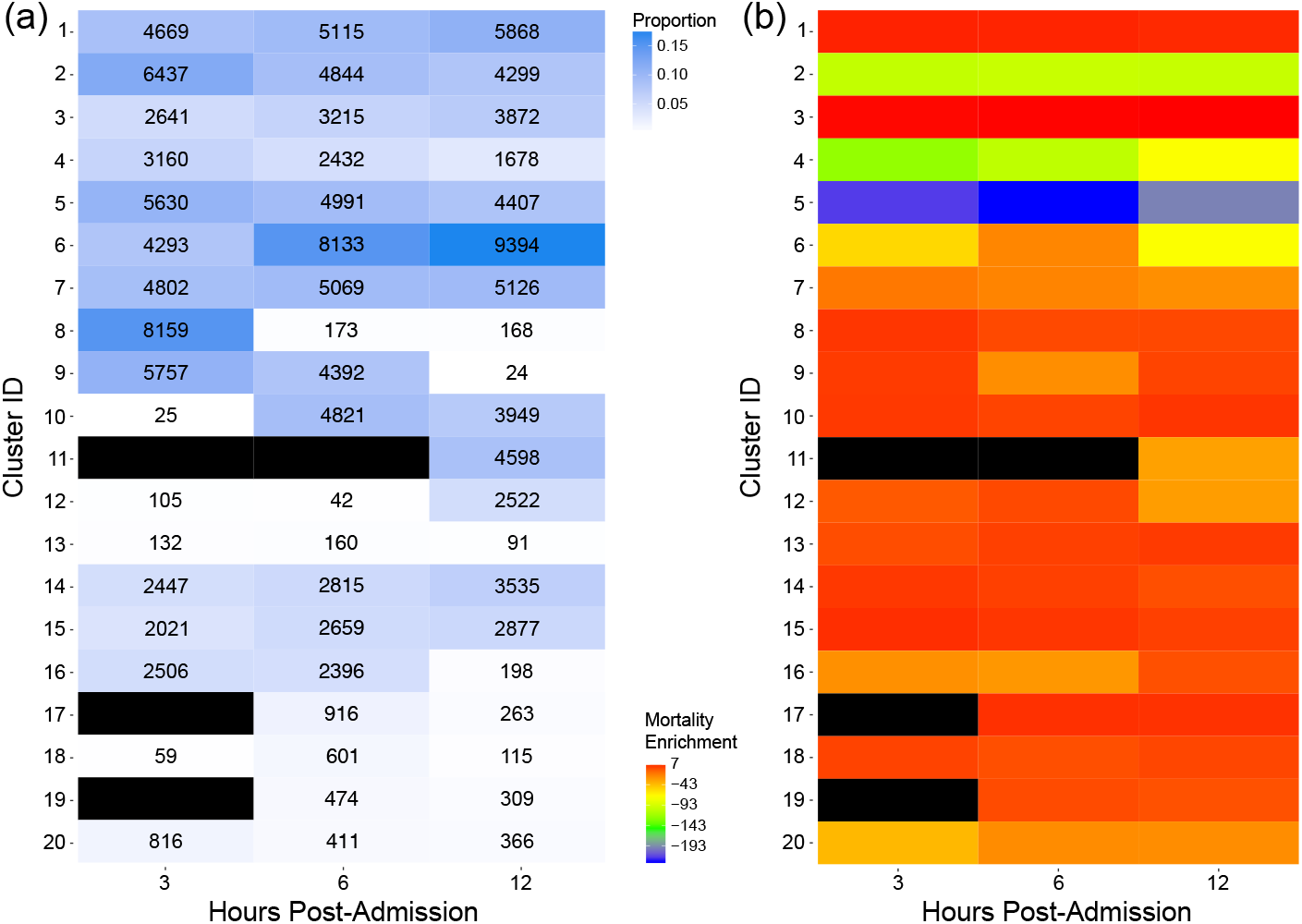
(a) Proportions of episodes assigned to each of the final 20 clusters for each post-admission period. The corresponding number of episodes assigned to each cluster are shown in each cell. (b) Mortality enrichment for each cluster and each post-admission period. The enrichment value is log(-log *(p)*) (for exposition purposes) where *p* is a one-tailed Fisher’s exact test p-value evaluating the significance of enrichment of mortality events for each cluster during a given post-admission period. Black cells indicate clusters to which no episodes were assigned during that post-admission period.

To further analyze temporal patterns in cluster membership, we then considered the full sequence of cluster assignments for each episode’s observations (*cluster trajectory*). This analysis resulted in 1,993 unique cluster trajectories, 18 of which were highly enriched for mortality events (Figure 5). The trajectory most enriched for mortality events involved episodes in which the patient remained in cluster 3 while other trajectories involved association with different clusters over the 12-hour post-admission period (e.g. trajectory 11 in Figure 5 starts in cluster 8 at 3 hours post-admission then transitions to clusters 6 and 3 at 6 and 12 hours post-admission, respectively). For these mortality-enriched trajectories, episodes tended to associate with a diverse set of clusters at 3 hours post-admission but generally transitioned into one of three clusters (1, 3, or 10) at 12 hours post-admission.

**Figure 5:**
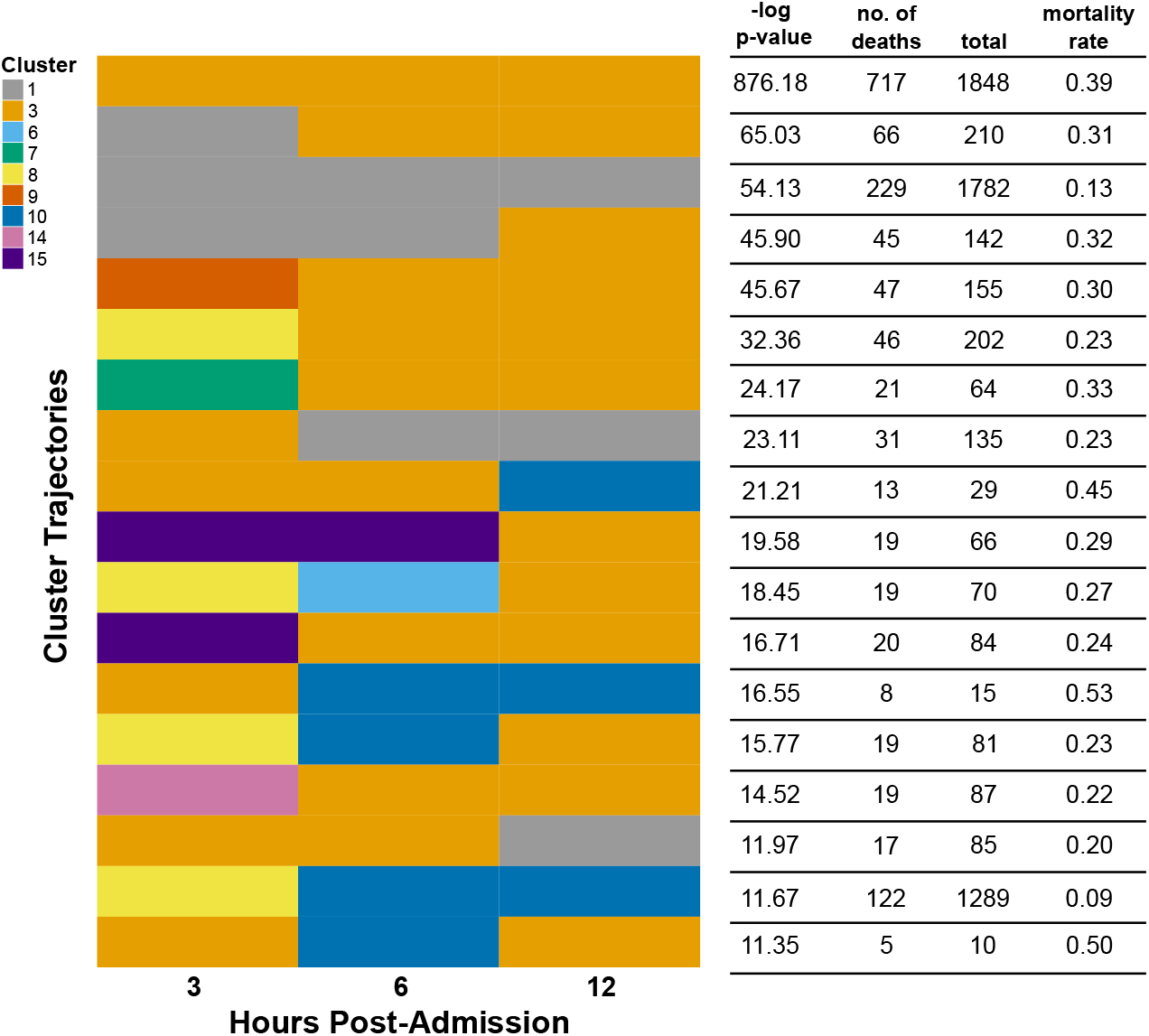
Cluster trajectories in the KPNC sepsis cohort enriched for mortality events. Trajectories are ranked in descending order by the -log p-value of a one-tailed Fisher’s exact test. A cluster trajectory appears in each row and consists of three cluster assignments, one for each of the three post-admission periods (3h,6h,12h). For example, the top cluster tra jectory indicates episodes that were assigned to cluster 3 at 3, 6, and 12 hours post-admission.

We assessed the physiological signatures of these three clusters at 12 hours post-admission by determining whether patient episodes associated with these clusters had features that were significantly different from those of the overall population (Figure 6; described in Methods, section 2.4). Interestingly, we find that while all three clusters associated with an elevated mortality risk, they reflect distinctly different physiological sub-populations, again highlighting the heterogeneity inherent in the presentation of sepsis. For example, while cluster 3 episodes showed significantly higher acute disease burden (LAPS2) than those episodes in clusters 1 and 10, cluster 1 episodes showed significantly higher chronic disease burden (COPS2). Notably, cluster 10 episodes showed significantly less chronic and acute disease burden than the overall population.

**Figure 6:**
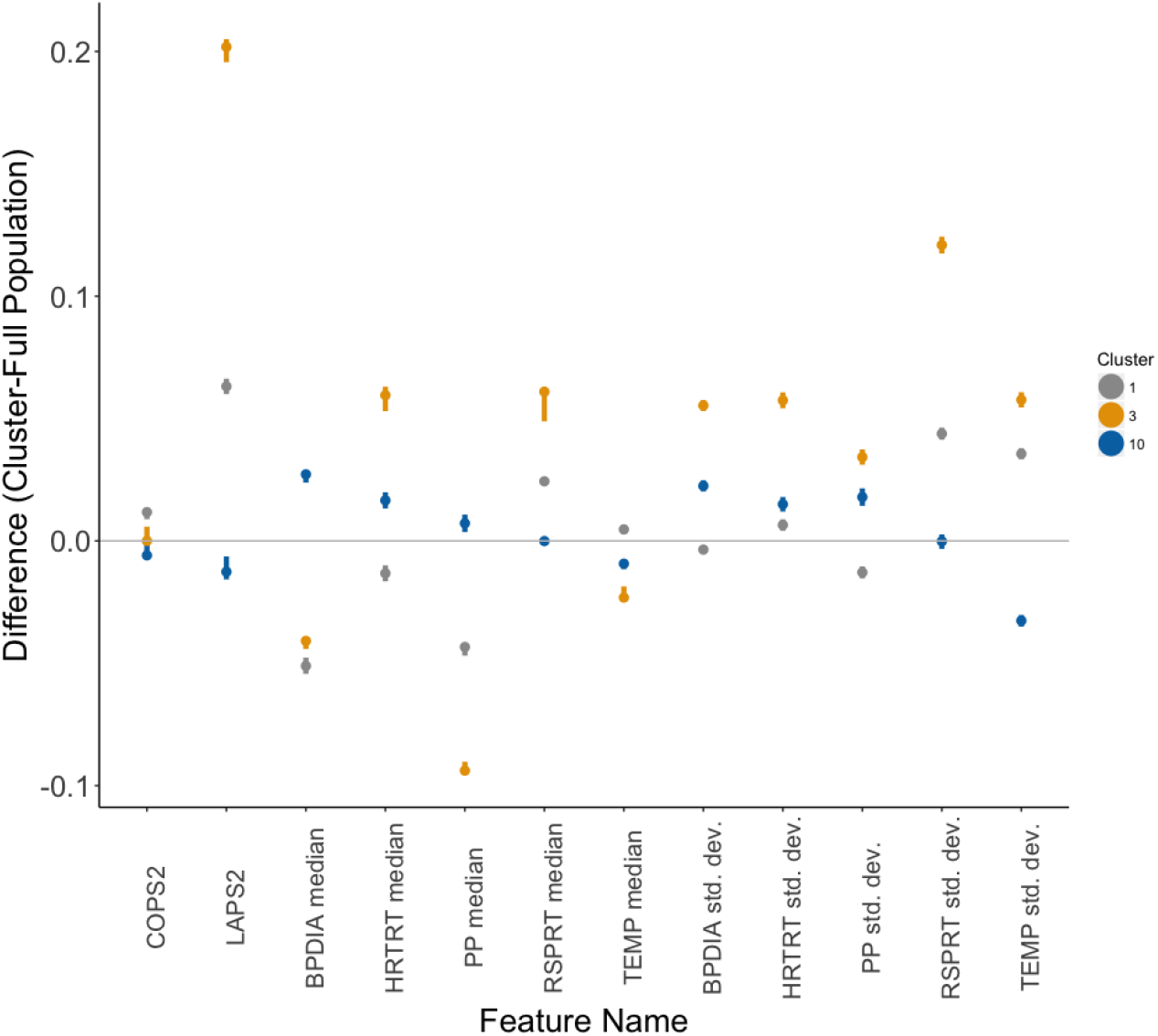
Estimated differences between cluster-specific and population values of each feature. The dots represent the estimated difference while whiskers represent the 95% confidence interval as computed by Wilcoxon rank-sum tests. The colors of each cluster’s dot and whiskers correspond to those in Figure 5. The lack of an overlap between these intervals and a difference of 0 (gray line) reflects a significant difference in cluster-specific values of the feature as compared to the overall population.

In addition, the vital sign characteristics of the different mortality-enriched cluster episodes were distinctly different. Vital sign median levels and standard deviations for cluster 3 generally deviated more substantially from those in the overall dataset when compared with the same characteristics in clusters 1 and 10. In particular, cluster 3 episodes showed significantly lower median blood pressure, pulse pressure, and body temperature, higher median heart and respiratory rates, and the highest vital sign variability. Cluster 1 vital sign patterns were generally similar to those of cluster 3. However, cluster 1 episodes did show significantly lower heart rate, higher body temperature, and generally lower vital sign variability for blood pressure, heart rate and pulse pressure. Cluster 10 episodes also showed different physiological trends: higher median blood pressure and pulse pressure, lower variability in body temperature, and respiratory rate median levels and variability close to those of the overall population. This analysis suggests that while a patient may be placed in a high-risk strata, their physiological state can be quite distinct from that of another high-risk patient. To validate the clinical interpretability of these physiological signatures, we provided them to our clinical collaborators with some orientation as to what was plotted to see if they could assign a coarse categorization to the clusters. They determined that cluster 3 episodes represented ’severe physiologic instability’, cluster 1 episodes indicated ‘respiratory failure in chronic disease’ and cluster 10 episodes represented ’moderate hemodynamic compromise’.

### 4.2. CMMs Enable Cluster-Based Visualization of Physiological Trends of Mortality Risk

In the previous analysis, we showed that the CMM can stratify populations into physiologically distinct subgroups that could highlight the need for different treatment regimens. To display these patterns, we adapted a technique often used in conjunction with random forest models: marginal importance plots [5, 21]. These plots show the expected mortality rate and 95% confidence intervals, at different values in the range of a given vital sign feature of interest (Figures 7a, 7b, 7c, and 7d; described in Figure 3). For this analysis, we fit the CMM model to the 12h dataset only and find trends largely reflecting current clinical knowledge and practices: median diastolic blood pressure and body temperature levels associated with low levels of mortality at 12 hours postadmission are within a range (i.e. “within normal limits”) outside of which the mortality rate begins to increase. The added advantage of this approach is 1) statistical assessment of confidence in the smoothed mortality rate estimates, 2) increases in dynamic range of mortality risk due to averaging over the different clusters in the CMM and 3) evaluation of nonlinear relationships between vital sign features and mortality risk.

In the case of median diastolic blood pressure (Figure 7a), the mortality rate varied between ∼6% (similar to the overall population rate) at values near 75 to rates in excess of 50% at values near 25, reflecting the importance of blood pressure management in sepsis ([14] and references therein). Likewise, in the case of body temperature (Figure 7c), mortality rates were at their lowest (∼5%) at approximately 98° F, climbing to rates of nearly 60% as body temperature dropped to 85° F. In the case of standard deviations of the different vital sign features (Figures 7b and 7d), the plots indicate that, in general, increased variability in a patient’s vital signs is associated with higher rates of mortality. However, this relationship was not linear (e.g. Figure 7d), suggesting that some variability was not harmful for patients, within a specific range dependent on each variable. Overall, in stratifying the population (as opposed to treating it as a homogeneous group) the CMM helps to identify “pockets” of demographic/physiological space that are strongly associated with increased mortality, thus facilitating more precise characterization of the marginal physiological trends associated with increased rates of mortality.

**Figure 7:**
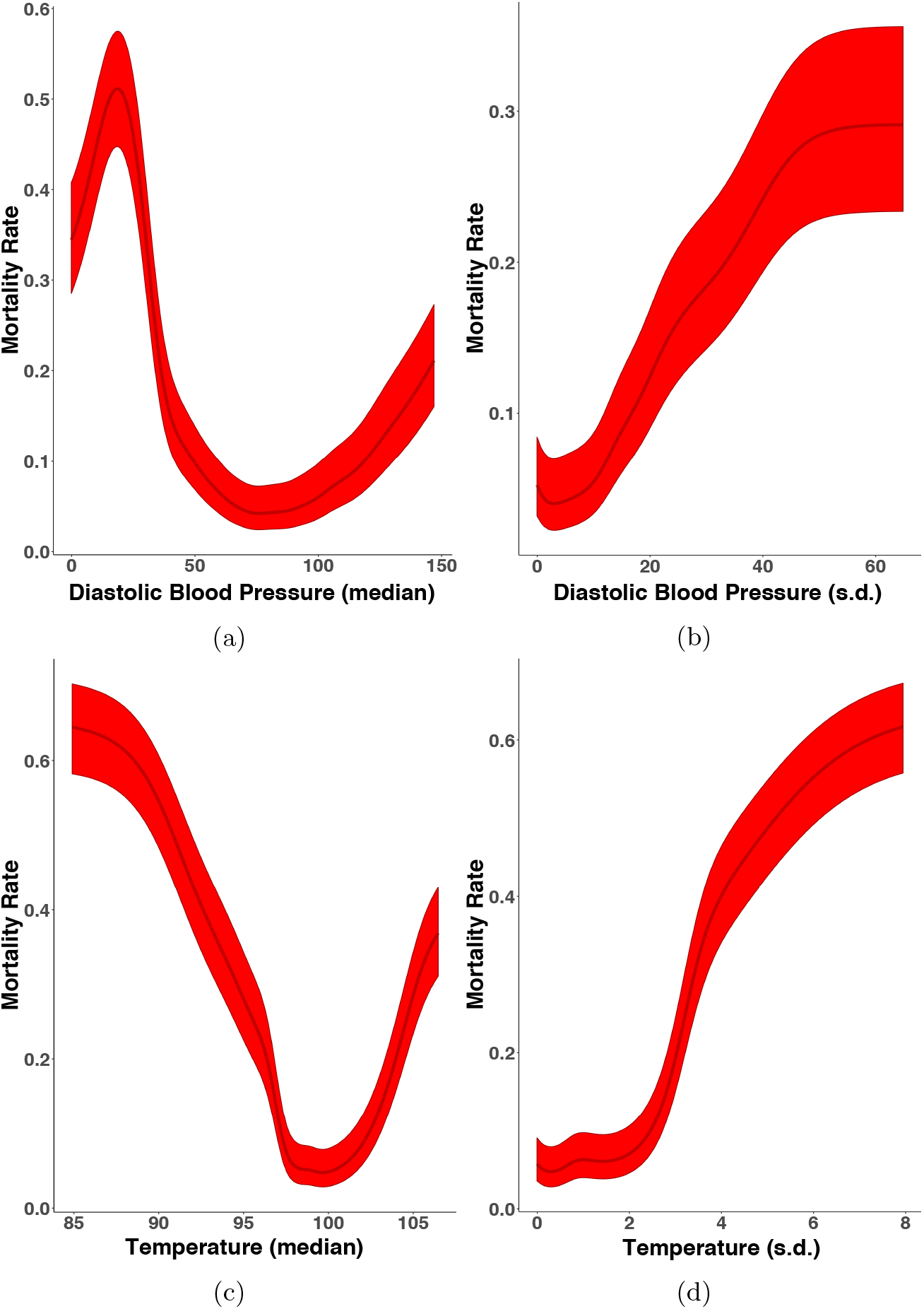
Marginal importance plots for diastolic blood pressure (median (a) and standard deviation (b)) and pulse pressure (median (c) and standard deviation (d)) at 12 hours post-admission. The dark line in the center of the band is the estimated mortality rate at each value of the vitals feature of interest while the lighter bands are 95% Wilson’s score intervals.

### 4.3. CMMs Provide Competitive Missing Data Imputation by Joint Learning of Feature Dependencies

As the CMM is a joint probability model, we can use the learned structure from a trained CMM to impute missing observations, a common task in EMR analysis. Indeed, recent unsupervised structure learning approaches have proven successful at this task [3]. We compare the CMM’s performance on a modified version of the 6h dataset (with 5% missing data introduced; see Methods) with that of three other imputation approaches: 1) MICE [6], a gold standard method for missing data imputation, 2) *k*-nearest neighbors, a nonparametric approach to imputation and 3) imputation with the population mean (our baseline). As MICE (based on fully conditional regressions) and CMMs (based on unsupervised learning) are designed for very different purposes, our goal in evaluating the CMM’s data imputation performance is to not to draw direct comparisons to stand-alone imputation methods, but rather to assess the situations in which CMMs might prove useful for data imputation. Figure 8 shows the distances, relative to population mean imputation, for select features using various imputation methods. Overall, the average relative distances across all features are 0.570 ± 0.001 (CMM using complete records), 0.524 ± 0.001 (CMM using population mean imputation), 0.398 ± 0.009 (MICE), 0.528 (5-nearest neighbors), and 0.501 (20-nearest neighbors), indicating the CMM’s general utility for missing data imputation.

**Figure 8:**
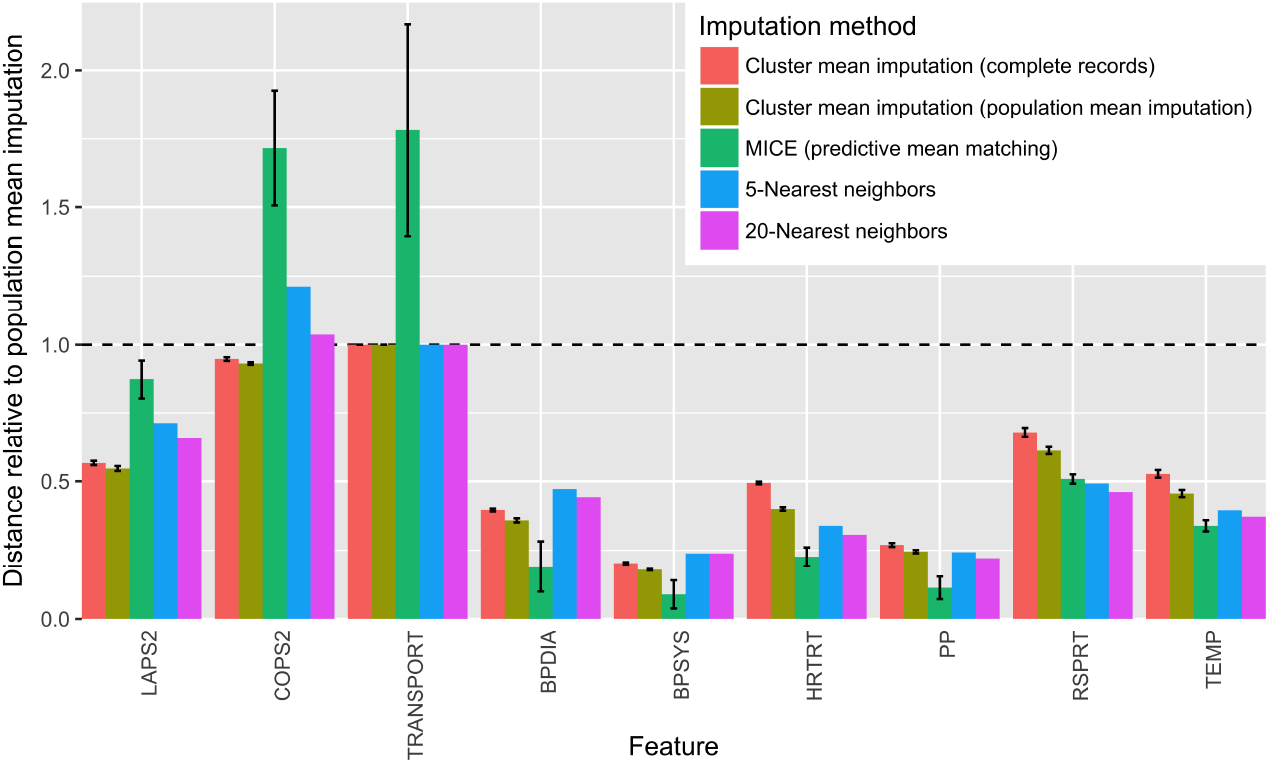
Distances between imputed and observed values from four different imputation methods for select features, relative to distances from population mean imputation (*D*_normalized_ = *D*_imp_. /*D*_pop. mean imp._), where *D* is the appropriate distance for each feature as described in section 3.6). Error bars represent standard deviation across imputations on 3 independent CMM fits (for CMM imputations) or across 20 independent imputations (for MICE). Performance was similar for TRANSPORT and two other categorical features (membership and facility codes) not shown. TRANSPORT, patient transport code; BPDIA, median diastolic blood pressure; BPSYS, median systolic blood pressure; HRTRT, median heart rate; PP, median pulse pressure; RSPRT, median respiratory rate; TEMP, median temperature.

For all features, CMM imputation using estimates based on population mean imputation outperformed CMM imputation using estimates based solely on complete records (Figure 8). This result was expected, as the former has over four times as much training data as the latter. For all summary statistics of vital signs, MICE outperformed both CMM imputation methods and was comparable in performance to k-nearest neighbors. As MICE leverages fully conditional data in its imputation and vitals are expected to be globally predictive of one another, this result is not surprising. No approach significantly improved over population imputation for the three categorical features in the dataset (membership status, transport code, and facility code).

CMM-based imputation did result in substantially smaller distances for one important physiological feature: LAPS2. While LAPS2—which is based on laboratory, vital sign, and neurologic information obtained within the 72 hours preceding hospitalization—shows very little conditional dependence on the other features in the dataset across the entire population (hence the slight improvement MICE yields over population mean imputation), the CMM seems to detect latent structure according to this measure of acute illness, offering a significant improvement over population mean imputation. This result corroborates our previous findings of population structure according to mortality risk.

## 5. Discussion & Conclusions

We demonstrated the flexibility and efficacy of our composite mixture model approach in analyzing multi-typed EMR datasets characterized by population heterogeneity and data missingness. The model can be used to stratify clinical populations according to risk of different unfavorable outcomes at different points in a hospitalization and to evaluate the marginal associations of patient physiology with higher rates of such outcomes. With the wide range of common EMR data types supported by the CMM, specification of a new model or extension of existing models to new data types is straightforward without requiring significant development efforts. Coupled with the cluster-based techniques we present here, the CMM can be used to statistically annotate groups of patients in an unbiased manner, an approach that can be easily be deployed as part of a patient monitoring system in other healthcare centers. Once the CMM is fit on episode data across a range of post-admission time periods, new patient feature vectors or patient feature vectors updated with new information and be fed to the model to determine changes in the probability of assignment of the patient to high-risk clusters. As a joint rather than conditional probability model, the CMM not only provides a useful framework to impute missing observations but also to predict outcomes of interest for held-out samples, enabling analyses that can complement purely predictive modeling studies.

Our work is most closely related to the unsupervised learning study Pivovarov et al. [30] and the supervised learning works by Halpern et al. [15] and Joshi et al. [19]. In [30], the authors transform different EMR data types (including clinical notes and ICD9 codes) into a single type of representation (bag-of-words) meant to be fit by a latent Dirichlet allocation-based mixture model [4]. In contrast, we directly model demographic and physiological variables of both discrete and continuous types from in-patient episodes.

Another important difference between this study and others is the data-driven approach we take to annotate and evaluate our clusters. Benchmarking of clustering approaches is problematic. In Halpern et al. [15] and [19], the authors take a supervised (or weakly supervised) approach to address this issue using “gold standard” (or “silver standard”) labels, evaluating the ability of their methods to recover manually curated phenotypes usually associated with certain clinical conditions (*external* evaluation; [12]). As we do not have access to such supervised information for sepsis [1, 8, 24] and are working in a more specific population of interest (e.g. suspected septic patients as opposed to all emergency department patients), we instead opt for an indirect evaluation of our clusters [12]: assessing their utility for the chosen goal of enriching patients into subgroups with statistically significantly higher rates of adverse health outcomes like mortality during hospitalization. We take this unbiased approach to discover and evaluate potentially uncharted, latent phenotypes relevant to sepsis treatment since manually curated conditions or prediction targets might introduce subjective bias into the kinds of phenotypes discovered. Consequently, an advantage of the CMM model, coupled with our statistically driven evaluation of the physiological signatures of high risk clusters, is its ability to generate clinically interpretable readouts of different patient subgroups of interest. While we make no claim of superiority in our clustering, we do believe that the flexibility and portability of the CMM, coupled with the novel cluster-based analyses we present here, provide a complementary, data-driven approach to clinical phenotyping and risk analysis.

The CMM is not without its drawbacks. For one, manually specifying the model template, or list of univariate distributions that model each feature column of a dataset, can be prohibitive in high feature dimensions. Also, while model fitting routines have been vectorized and the component distributions (i.e. exponential family distributions) are amenable to embarrassing data parallelism [28], our approach was not optimized further. As such, estimation of a CMM can be computationally intensive (e.g. approximately 3.5 hours on a single core for the 3 hr dataset, with 53,659 episodes and 32 features, fitting to 250 (the optimal number) clusters). In previous non-medical applications, we overcame this computational burden by leveraging scalable high-performance computing software framework for streaming data analysis [37]. We are currently extending our model fitting software to automatically identify “default” distributions for a given feature column as well as exploring scalable (i.e. data-parallel) estimation routines.

In addition to demographic information, we opted to model a particular set of patient vital sign statistics representative of level and variability over fixed post-admission periods. However, this approach ignores dynamic patterns in the vital signs including changes in the cross-correlation between different vital sign types that might also be indicative of changes in patient health [27] and could be included as features. In addition, an alternative approach to modeling the vital signs would be to extend the CMM with distributions appropriate for time series data (e.g. a Gaussian process; [31, 22]). Such an extension could potentially reduce the number of parameters in the overall CMM and lessen model specification and estimation time. Moreover, while not included in our analyses, lab test results can aid in the risk stratification of patients and, like vital signs, provide clinically actionable targets for intervention to improve patient health. As part of our ongoing and future work, we are investigating these extensions to our analyses.

We achieved competitive missing data imputation performance with the CMM, though it is important to note how that analysis also highlights differences between unsupervised and supervised learning approaches. Regression-based methods like MICE make predictions based on population-wide conditional information. Thus, features for which MICE performs well could indicate strong conditional dependencies between those features and all other features at the population level. In contrast, CMM-based imputation makes predictions based on identifying latent local structure, identifying subpopulations of the full dataset that are more similar to one another. In particular, the CMM uncovers latent structure with respect to features like LAPS2, leading to improved imputation performance, even though the current formulation of the model does not include conditional dependencies. It is worth noting that our assumption of conditional independence among feature dimensions given a mixture component represent one potential model structure of the CMM. Conditional dependencies can be introduced into the CMM rather straightforwardly (e.g. replacing univariate normal distributions with linear regressions conditioned on other features) and will be the subject of future work. Taken together, these results suggest that the CMM could be used in concert with other data imputation approaches to address missing data challenges on a per-feature basis.

Our CMM-based analysis and data-driven cluster annotation of the KPNC sepsis cohort revealed physiologically distinct subpopulations associated with elevated rates of mortality. In a patient monitoring context, a patient’s observation vector could be passed as input to our fitted model at different points in their hospitalization to evaluate the probability of their association with high-risk clusters. Moreover, an important follow-up analysis could identify patterns in those episodes assigned to high risk clusters at 3 hours post-admission that were assigned to low risk clusters later in their hospitalization (and vice versa). Specifically, as our full dataset also includes discrete time series of the medications and procedures ordered by the clinician during each hospitalization, we can evaluate whether certain medications or procedures were over-represented in trajectories in which patients transitioned into clusters of higher or lower risk. Overall, our CMM framework provides a valuable decision support tool to characterize and stratify heterogeneous clinical populations, capturing clinically relevant changes in patient physiology while accounting for the data missingness and wide variety of data types inherent in large-scale electronic medical record data.

## 6. Acknowledgements

The authors would like to thank Priyadip Ray and Tim Sweeney for comments on the manuscript. Dr. Vincent X. Liu is funded by NIH grant (NIH K23GM112018). This work was performed under the auspices of the U.S. Department of Energy by Lawrence Livermore National Laboratory under contract DE-AC52-07NA27344 (LLNL-JRNL-730845-DRAFT).

